# Microinjection-based Single-Cell Toxicological Assessment Reveals How Physiological Levels of PFOS Impair Oocyte Maturation and Developmental Competence

**DOI:** 10.64898/2026.05.26.727938

**Authors:** Hasanur Alam, Shuangqi Wang, Juan Dong, Vidhi Patel, Wenjie Yang, Leyi Wang, Huanyu Qiao

## Abstract

Perfluorooctanesulfonic acid (PFOS) is a persistent environmental contaminant widely detected in human serum and follicular fluid and has been associated with reduced implantation rates and female fertility. However, its direct effects on mammalian oocyte maturation remain poorly understood. Here, we developed a microinjection-based single-oocyte toxicological assay to directly evaluate how physiologically relevant PFOS concentrations affect mouse oocyte maturation and early embryonic development. Microinjection of PFOS at follicular-fluid level (5.6 nM) and occupational exposure level (60 nM) significantly reduced germinal vesicle breakdown (GVBD) and polar body extrusion (PBE) rates compared with water-injected controls. Notably, all tested concentrations (2.4 nM serum level, 5.6 nM, and 60 nM) induced abnormal polar-body formation, disrupted meiotic spindle morphology, and increased the proportion of unhealthy oocytes. PFOS exposure also significantly elevated intracellular reactive oxygen species (ROS) levels and mitochondrial membrane potential at 5.6 nM, indicating oxidative stress and mitochondrial dysfunction. Cytological analyses revealed chromosome misalignment and widened metaphase I plates, suggesting chromosome missegregation and subsequent prometaphase II arrest with defective polar bodies. Single-cell RNA sequencing of PFOS-treated oocytes exhibiting abnormal small polar bodies identified distinct transcriptional signatures, including dysregulation of genes involved in mRNA processing, chromosome segregation, mitochondrial function, and cell division. Functionally, these oocytes failed to progress beyond the 2-cell stage following in vitro fertilization, indicating loss of developmental competence. Collectively, these findings demonstrate that PFOS directly disrupts meiotic progression through spindle defects, oxidative stress, and transcriptional dysregulation, ultimately compromising oocyte quality even at environmentally relevant exposure levels.

**Environmental Implication:** PFOS is a persistent environmental contaminant widely detected in human serum and follicular fluid. Our findings demonstrate that PFOS at physiologically relevant levels can impair oocyte maturation, disrupt meiotic chromosome segregation, and compromise early embryonic development. By using a single-oocyte toxicological assay, we reveal that even low-dose PFOS exposure can induce oxidative stress and transcriptional dysregulation. These results highlight the potential reproductive risks of chronic PFOS exposure and underscore the importance of stricter environmental monitoring and regulation to protect female reproductive health and fertility. This novel assay also has the potential to redefine safety thresholds for other environmental toxicants.

## 1. Introduction

Infertility is defined as the failure to achieve pregnancy after at least 12 months of regular, unprotected sexual intercourse despite active attempts to conceive. Globally, approximately 8–12% of couples experience infertility (*1*). Increasingly, environmental pollutants have been recognized as key contributors to reproductive dysfunction and the rising infertility rates (*2*). Common environmental toxicants - such as endocrine disruptors, heavy metals, and pesticides - are pervasive in modern life and can impair gamete quality and reduce overall reproductive success. Although the effects of these toxicants are often subtle and cumulative, their long-term influence on fertility is substantial.

Per- and polyfluoroalkyl substances (PFAS) are a large class of environmentally persistent chemicals frequently detected in drinking water (*3*, *4*) and various consumer products, including water-resistant clothing (*5*), upholstered furniture (*6*), non-stick kitchenware (*7*), baking parchment (*8*), and food packaging (*9*). Epidemiological studies suggest that PFAS exposure can reduce fertility in women by up to 5-10% (*10*).

Among the PFAS chemicals, perfluorooctanesulfonic acid (PFOS) is of particular concern due to its surfactant properties, chemical stability, and strong bio-accumulative potential. With an estimated half-life of approximately 27 years (*11*), PFOS persists in the environment and accumulates in humans, livestock, and wildlife, raising serious concerns about its toxicity (*12–14*). Growing evidence links PFOS exposure to reproductive dysfunction, including endocrine disruption, impaired gametogenesis, and defective embryo development (*15*). In females, PFOS exposure has been associated with impaired folliculogenesis, reduced estradiol synthesis, and poor oocyte quality (*16*). Our previous study demonstrated that the addition of 600 µM PFOS to the culture medium impairs oocyte maturation (*17*). However, the exact amount of PFOS that enters oocytes during *in vitro* culture remains unknown and most previous studies employ PFOS concentration far exceeding those encountered in real-world human exposures. Physiologically relevant PFOS levels are approximately 2.4 nM in human serum and 5.6 nM in follicular fluid levels, while occupationally exposed individuals may reach levels up to 60 nM(*18–20*). The marked discrepancy between measured physiological levels and experimentally applied concentrations limits the ability to accurately assess PFOS toxicity in real world. Therefore, it is essential to examine PFOS effects at both physiologically and occupationally relevant exposure levels.

Given the reproductive risks associated with PFOS, it is particularly important to understand how low-dose exposure affects oocyte maturation—a tightly regulated process required for successful fertilization and embryo development. Oocyte maturation proceeds from the germinal vesicle (GV) stage to metaphase II (MII) stage and make oocytes competent for fertilization (*21*). During meiosis I, homologous chromosomes must align precisely at the metaphase plate to ensure correct kinetochore-microtubule attachments and achieve accurate segregation (*22*). Unlike mitotic cells, oocytes lack centrosomes and instead rely on multiple microtubule organizing centers (MTOCs) to assemble the meiotic spindle, rendering the process inherently more error-prone (*23*). Errors in chromosome alignment – arising from defects in kinetochore–microtubule interactions or spindle architecture - can lead to lagging chromosomes, aneuploidy, and meiotic arrest (*24*). Even mild disturbances, such as oxidative stress or chemical exposure, can disrupt chromosome alignment and compromise oocyte quality (*25*, *26*). PFOS has been shown to elevate intracellular reactive oxygen species (ROS) levels, which leads to oxidative stress, alter mitochondrial membrane potential, induce DNA damage, and trigger apoptosis, all of which compromising oocyte quality and developmental competence (*17*). However, the effects of PFOS at physiologically relevant concentrations on meiotic progression remain poorly understood.

Microinjection offers a precise approach for assessing toxicant effects by bypassing physiological metabolism, penetrating the oocyte membrane barrier, and delivering defined intracellular doses. In this study, we developed a microinjection-based single-oocyte toxicological assay by introducing PFOS into mouse oocytes at concentrations reflecting physiological and occupational exposure levels. We examined key meiotic events–including germinal-vesicle breakdown (GVBD), polar-body extrusion (PBE), spindle morphology, chromosome alignment, and mitochondrial function–to determine how low-dose PFOS perturbs oocyte maturation. To further explore the mechanisms underlying PFOS-induced dysfunction, we evaluated oxidative stress by measuring ROS production and mitochondrial membrane potential. In addition, PFOS-injected oocytes frequently extruded abnormally small polar bodies. By performing single-cell RNA sequencing on oocytes exhibiting normal versus small polar bodies, we characterized the transcriptomic differences associated with these phenotypes. Together, our findings provide new insights into the cellular and molecular consequences of PFOS exposure at environmentally relevant levels, highlighting its detrimental effects on oocyte meiotic competence and transcriptional regulation.

## 2 Materials and Methods

### 2.1 Chemicals

Perfluorooctanesulfonic acid (PFOS), a perfluoroalkyl sulfonic acid (PFSA), is from Synquest Laboratories (Alachua, FL, USA). Unless otherwise specified, all other chemicals were purchased from Sigma-Aldrich (St. Louis, MO, USA).

### 2.2 Animals

CD-1 mice (Charles River Laboratories, Wilmington, MA) were used in the study. They were housed in the Animal Care Facility at the University of Illinois Urbana-Champaign (UIUC) under a 12-hour dark/ 12-hour light cycle at 22±1°C. Food and water were provided *ad libitum*. The UIUC Institutional Animal Care and Use Committee (IACUC) approved animal handling and procedures.

### 2.3 Cumulus-oocyte-complexes (COCs) collection and *in vitro* maturation

Female mice (4-6 weeks old) were euthanized and dissected for ovary collection. The ovaries were washed in PBS, and any excess fat and surrounding tissues were removed at this time. The ovaries were then transferred to pre-warmed (37°C) M2 media. A sterile tweezer was used to hold the ovary, and a syringe needle was used to rupture the antral ovarian follicles, releasing COCs. The COCs were collected into a separate droplet of pre-warmed M2 media, microinjected with PFOS, and washed three times. The washed COCs were then transferred to pre-warmed *in vitro* maturation (IVM) medium covered with mineral oil. The IVM medium consisted of minimum essential medium Eagle-alpha (α-MEM) supplemented with 10% fetal bovine serum (FBS), 200 mIU follicle-stimulating factor (FSH), and 50 µg/mL sodium pyruvate. The COCs were cultured at 37°C in 5% CO_2_ for 6 hours. Following incubation, cumulus cells were removed by repeated pipetting, and GVBD rates were examined. Denuded oocytes were then washed three times in pre-warmed M16 and transferred into a fresh droplet of M16 medium covered with mineral oil. The oocytes were further incubated at 37°C in 5% CO_2_ for 18 hours. At the end of the culture period, the rates of normal polar body extrusion, defective polar body formation, and unhealthy oocytes were examined. Oocytes with granular cytoplasm, cytoplasmic shrinkage, and detachment of the oocyte membrane from the zona pellucida (increase in gap of perivitelline space) were classified as unhealthy. Polar bodies exhibiting the absence of a spindle and reduced size were classified as defective.

### 2.4 Single-cell microinjections and culture (Figure S1)

Borosilicate glass capillaries (TW100-4, World Precision Instruments Inc, FL) were pulled into either holding or microinjection capillaries using a horizontal micropipette puller (P-97, Sutter Instrument Company, CA). The holding capillary held and stabilized the COCs during PFOS microinjection. Diluted PFOS was backfilled into the microinjection capillaries using a microloader 100 mm, 20 μL pipette tips (Fisher Scientific, MA). The microinjection capillary was attached to the micropipette holder of a hydraulic pump and positioned on a micromanipulator. The opposite end of the micropipette holder was connected to a Femtojet4i (Calibre Scientific, OH), which regulated injection pressure and duration to ensure precise PFOS delivery into oocytes.

To determine PFOS injection volumes, we measured the diameter of the expelled PFOS droplet in mineral oil and calculated its volume using the formula V = 4/3 πr³. Under the defined injection parameters (50 hPA injection pressure, 0.10 s injection time, and 20 hPA compensation pressure), the PFOS droplet volume is 0.523 pL. The average oocyte diameter is 80 µm, and its volume was calculated as 268.083 pL using the same formula. To achieve the three target final PFOS concentrations within oocytes 2.4 nM (blood serum level), 5.6 nM (follicular fluid level), and 60 nM (occupational exposure level), the required stock concentration in the microinjected PFOS droplet was determined using this formula, X = 268.083×Oocyte PFOS Concentration/0.523. The corresponding stock concentrations were 1230.21 nM, 2870.49 nM, and 30755.22 nM, respectively. Control oocytes received an equivalent volume (0.523 pL) of sterile water under identical conditions.

The micromanipulator guided the microinjection capillary against the holding capillary for precise penetration through the zona pellucida. PFOS was injected into the oocyte cytoplasm by controlling pressure and duration. All COCs underwent the same procedure and were subsequently cultured in α-MEM medium supplemented with 10% FBS, 200 mIU FSH, and 50 µg/mL sodium pyruvate at 37°C in a 5% CO_2_ for 6 hours, allowing recovery from injection shock. After this period, cumulus cells were removed, and denuded oocytes were cultured in M16 medium at 37°C in a 5% CO_2_ incubator for an additional 18 hours.

### 2.5 Immunofluorescent staining

Oocytes were fixed in phosphate-buffered saline (PBS, pH 7.4) containing 4% paraformaldehyde for 30 minutes at room temperature. After fixation, they were washed three times with a washing buffer (0.1% tween-20 and 0.01% Triton X-100 in PBS) before being permeabilized in 0.5% triton X-100 in PBS for 20 minutes at room temperature. After three additional washes, the oocytes were blocked in 3% bovine serum albumin (BSA) for 1 hour at room temperature. They were then washed three times and incubated with FITC-labelled mouse anti-α-tubulin (1:200) for 1 hour at 37°C in a 5% CO_2_ incubator. To visualize F-actin, oocytes were treated for 5 minutes at room temperature with Alexa Fluor 594 Phalloidin (5 U/mL; Life Technologies) diluted in PBS. After three washes, their DNA was counterstained with 1 ug/ml of 4’-6-dimidine-2’-phenylindole dihydrochloride (DAPI) for 10 minutes at room temperature. Subsequently, the oocytes were washed twice and imaged using a Nikon A1R confocal microscope (Nikon Instruments Inc., NY). The images were processed using NIS-Elements software.

### 2.6 Measurement of intracellular reactive oxygen species (ROS) levels

Intracellular ROS level in control and PFOS-injected oocytes was assessed using an oxidation-sensitive fluorescent probe, 2′,7′-dichloro-dihydrofluorescein diacetate (DCFH-DA). Briefly, oocytes were incubated in M2 medium with 5 µM DCFH-DA at 37°C in a 5% CO_2_ incubator for 30 min. After incubation, oocytes were washed twice in M2 medium to remove excess DCFH-DA. Once inside the oocyte, the diacetate groups of DCFH-D are cleaved by cellular esterases, converting DCFH-DA into the non-fluorescent intermediate DCFH, which is further oxidized by ROS to yield the fluorescent product DCF (2′,7′-dichlorofluorescein). Green fluorescent signals of DCF were immediately imaged by a Keyence BZ-X fluorescence microscope (Keyence Corporation, IL). All images were acquired under identical exposure and acquisition settings for both control and PFOS-injected oocytes. Signal intensity was quantified by measuring the intensity within a defined region of interest (ROI) using Image J software (National Institutes of Health, USA).

### 2.7 Measurement of mitochondrial membrane potential (MMP)

The JC-1 mitochondrial membrane potential detection kit was used to measure mitochondrial membrane potential in oocytes. PFOS was microinjected into oocytes within COCs, which were then cultured at 37°C in a 5% CO_2_ incubator for 2 hours. After culture, cumulus cells were removed, and oocytes were incubated in 1X JC-1 solution (Biotum, Inc., CA) at 37 °C in a 5% CO_2_ incubator for 30 minutes in the dark. Oocytes were then washed 2-3 times in M2 medium, and the fluorescence signal was immediately examined by Keyence BZ-X microscope (Keyence Corporation, IL). Images of control and PFOS-treated oocytes were taken using identical imaging parameters. Signal intensity was quantified by measuring the intensity of the ROI using Image J software (National Institutes of Health, USA).

### 2.8 *In vitro* fertilization of oocytes

Male mice older than 2 months were euthanized, and the cauda epididymides were carefully dissected in pre-warmed M2 medium. Each cauda was placed in a 400 µL drop of EmbryoMax Human Tubal Fluid (HTF) medium under mineral oil. A 16-gauge needle was used to incise the cauda and release sperm from the cauda epididymides. The released sperm were incubated at 37 °C in a humidified atmosphere containing 5% COL for 5 minutes, after which residual tissue fragments were removed. The sperm suspension was then incubated for an additional 1 hour at 37 °C under 5% COL to allow capacitation. *In vitro* matured oocytes were transferred into a 50 µL drop of HTF medium. The sperm suspension was diluted 1:2 with an additional 400 µL of HTF. A 10 µL aliquot of diluted sperm suspension was gently added to the oocyte-containing drop. Co-incubation was carried out at 37 °C in a 5% COL incubator for 4 hours. After fertilization, excess sperm attached to the oocyte were removed by gentle pipetting. Presumptive zygotes were transferred to a fresh 50 µL drop of EmbryoMax CZB medium in a 35 mm culture dish and cultured at 37 °C in a 5% COL incubator for 48 hours.

### 2.9 Single-cell RNA sequencing

Single-cell RNA sequencing was performed using the Smart-seq3 protocol to investigate the transcriptomic profiles of MII stage oocytes derived from PFOS-injected oocytes. For each experimental group, one MII oocyte was collected into lysis buffer containing RNase inhibitor. mRNA was reverse-transcribed using an oligo(dT) primer carrying a 5′ anchor sequence. To enable full-length cDNA synthesis, a template-switching oligonucleotide (TSO) was included during the reverse transcription step. The resulting cDNA was then amplified via PCR and purified with magnetic beads. Quantification of amplified cDNA was performed using the Qubit™ dsDNA HS Assay Kit (Thermo Fisher), and fragment size distribution was evaluated using the Agilent 4200 TapeStation. Library construction was carried out with the Nextera XT DNA Library Prep Kit for Illumina according to the manufacturer’s instructions. The final libraries were assessed for quality and subsequently sequenced on the Illumina MiSeq platform using 150 bp paired-end reads (PE150). Raw reads were processed using Cutadapt (2.8) and Trimmomatic (0.39) to remove sequencing adapters and low-quality bases. High-quality reads were then mapped to the mouse genome (mm10) using Hisat2. Differential gene expression analysis was conducted using DESeq2 (1.38.3), with significantly differentially expressed genes (DEGs) defined by a |logL fold change| > 1 and p-value < 0.05. Functional enrichment analysis based on Gene Ontology (GO) terms was performed using the ClusterProfiler (4.6.2) package.

### 2.10 Statistical Analysis

Each treatment was performed at least three times. Data was presented as mean ± SEM. An unpaired two-tailed t-test was applied to analyze differences between the control and PFOS-injected groups. Statistical significance was set at \**P* < 0.05, \*\**P* < 0.01, and \*\*\**P* < 0.001. Graphs were made using OriginLab 2024 (Northampton, MA, USA).

## 3. Results

### 3.1 PFOS microinjection induces mouse oocyte maturation failure

To evaluate the effects of PFOS microinjection on mouse oocyte maturation, oocytes within COCs were microinjected with increasing concentrations of PFOS (2.4 nM, 5.6 nM, and 60 nM). 2.4 nM, 5.6 nM, and 60 nM correspond to blood serum, follicular levels, and occupational exposure levels. The GVBD rate was recorded after 6 hours of *in vitro* culture, while the PBE rate was assessed after 24 hours. As shown in Fig. 1A and 1B, PFOS microinjection at 5.6 nM and 60 nM had significantly reduced the GVBD rate compared to the water-injected control (*P* < 0.01 and *P* < 0.001, respectively). However, there was no significant difference in GVBD rate between 2.4 nM PFOS microinjection and the control. We further examined the effects of PFOS microinjection on PBE after 24 hours of culture. The PBE rates of mouse oocytes were significantly reduced in the 5.6 nM (47.05 ± 3.66%) and 60 nM (33.33 ± 9.62%) microinjection groups compared to the control (71.42 ± 1.58%) (*P* < 0.01 and P < 0.001, respectively; Fig. 1A and 1C). However, no significant difference was observed between the 2.4 nM PFOS microinjection and the control. PFOS microinjection significantly increased the proportion of unhealthy oocytes in the 2.4 nM group (*P* < 0.01), and more markedly in the 5.6 nM and 60 nM groups (*P* <0.001) compared to the control (Fig. 1D). Taken together, these observations revealed that PFOS microinjection blocked meiotic progression in mouse oocytes, which leads to the failure of oocyte maturation. Moreover, PFOS microinjection negatively affects the oocyte health and thereby increases the proportion of unhealthy oocytes.

**Figure 1.**
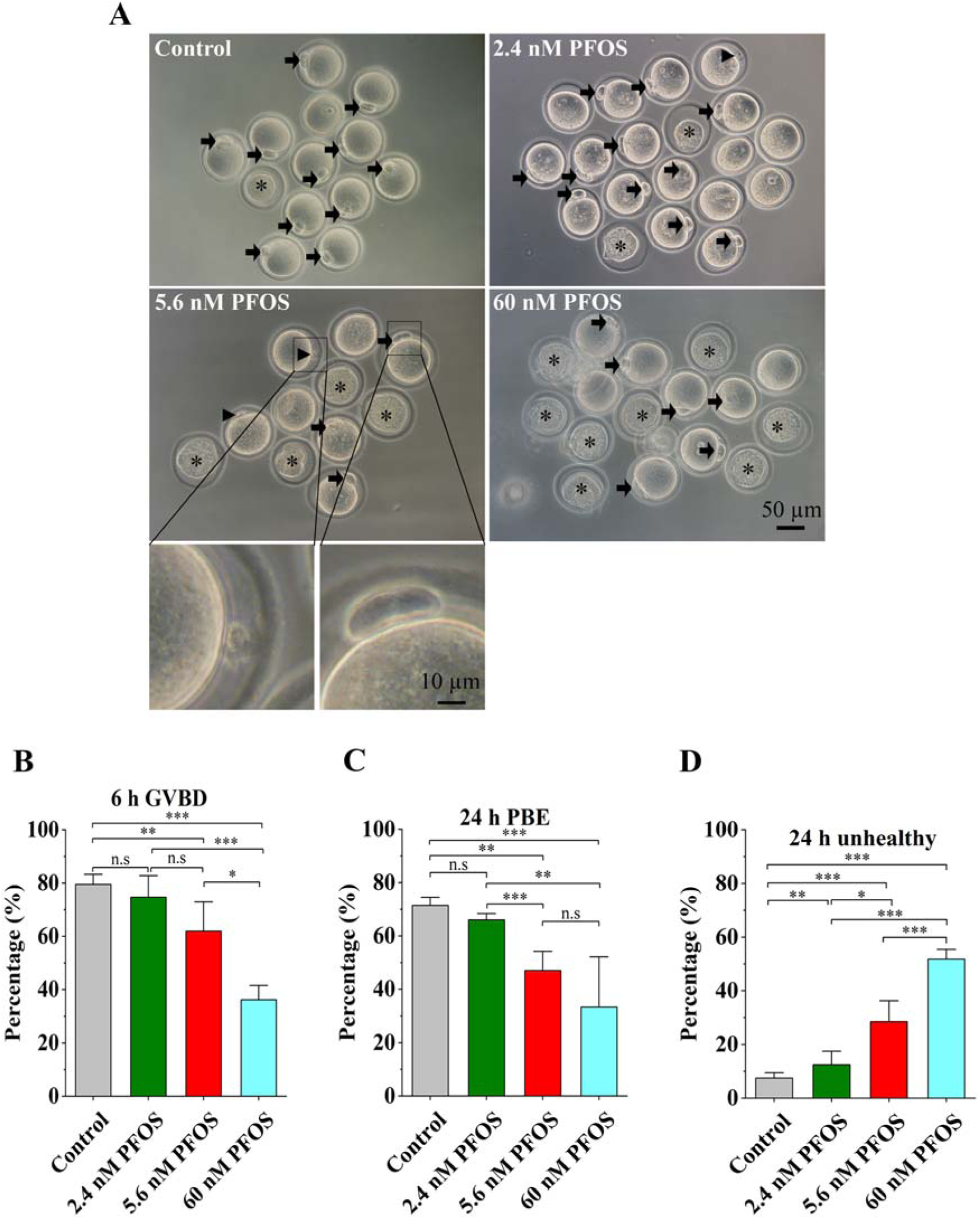
Effects of PFOS microinjection on mouse oocyte *in vitro* maturation. (A) Representative images showed the morphology of oocytes after 24 h *in vitro* culture in the control and PFOS-injected groups. The black arrows highlighted the oocytes with a normal polar body. The asterisk sign indicated oocytes with unhealthy conditions. Scale bar, 50 μm. (B) The rates of GVBD in the control and PFOS-injected groups after 6-hour culture. (C) The PBE rates in the control and PFOS-injected groups after 24-hour culture. (D) The unhealthy oocyte rates in the control and PFOS-injected groups after 24-hour culture. Data were presented as mean ± SEM of at least three independent experiments. \**P* < 0.05, \*\**P* < 0.01, and \*\*\**P* < 0.001 compared with control.

### 3.2 Effects of PFOS microinjection on the levels of reactive oxygen species (ROS) in mouse oocytes

Our previous studies have shown that the addition of 600 µM PFOS to the culture medium increases reactive oxygen species (ROS) levels of mouse oocytes, leading to DNA damage, apoptosis, and impaired oocyte quality (*17*). Thus, we hypothesized that PFOS microinjection at the concentrations corresponding to blood serum and follicular fluid levels would increase ROS levels in mouse oocytes, which may lead to the aforementioned detrimental effects. Our results showed that the fluorescence of DCF, a marker of intracellular ROS, was notably brighter in PFOS-microinjected oocytes than that in the control oocytes (Fig. 2A). Quantitative analysis revealed that ROS levels were significantly elevated in both 2.4 nM and 5.6 nM PFOS groups compared to the control group (*P* < 0.001, Fig. 2B). However, no significant difference was observed between the 2.4 nM and 5.6 nM groups, indicating that the microinjection of both blood serum and follicular-fluid levels of PFOS are sufficient to increase ROS levels in mouse oocytes.

**Figure 2.**
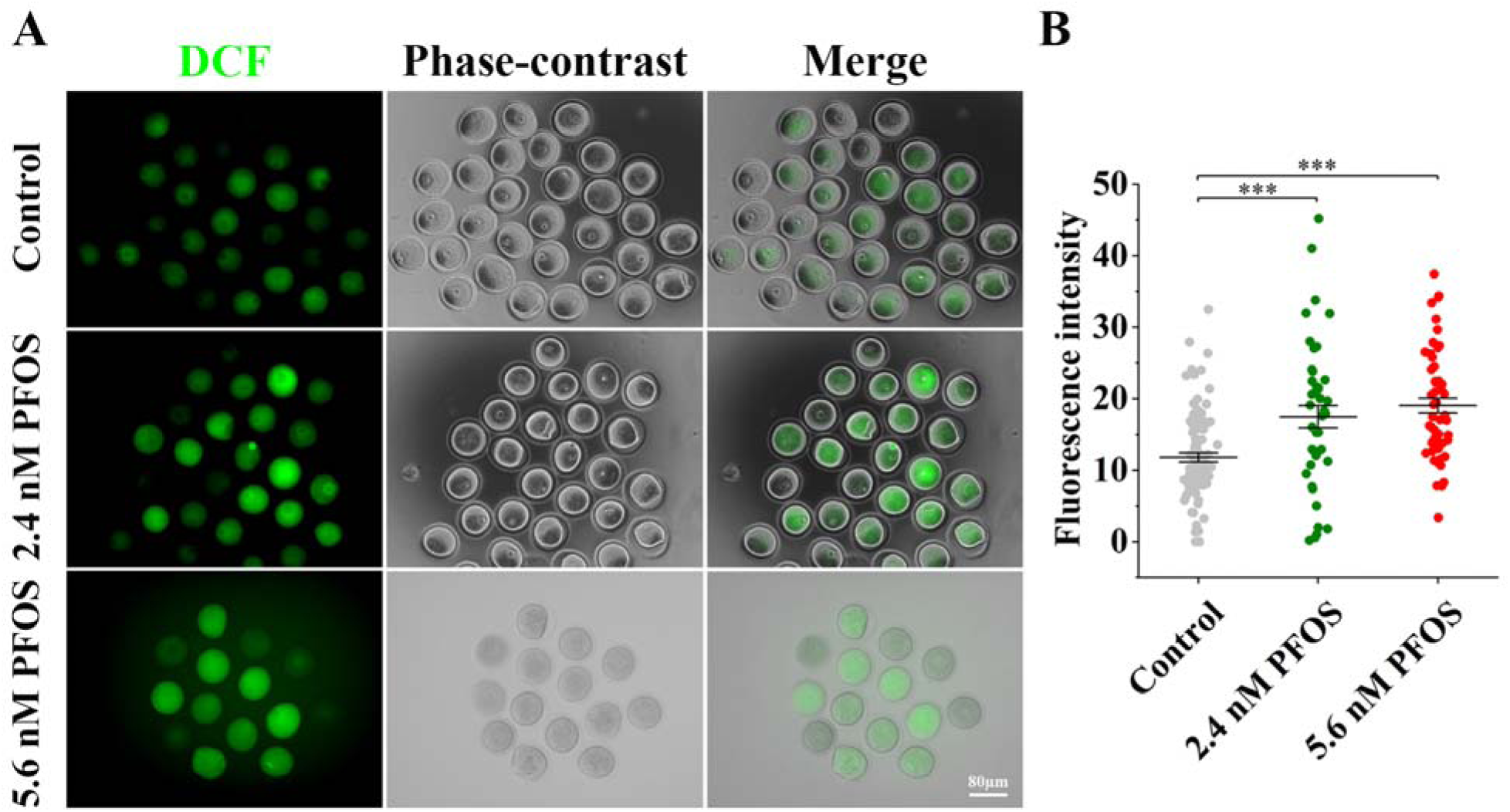
Effects of PFOS microinjection on ROS levels in mouse oocytes. (A) Representative images of DCF fluorescence (green) in water-injected control, 2.4 nM, and 5.6 nM PFOS-injected oocytes. Scale bar: 80 μm. (B) Quantitative analysis of DCF fluorescence intensity in the control, 2.4 nM, and 5.6 nM PFOS-injected groups. A total of 98 oocytes (control), 44 oocytes (2.4 nM PFOS), and 53 oocytes (5.6 nM PFOS) were analyzed for ROS levels. Data are presented as mean fluorescence intensity ± SEM from at least three independent experiments. t-test, \*\*\**P* < 0.001 versus control.

### 3.3 PFOS microinjection enhances oocyte mitochondrial membrane potential

To investigate mitochondrial function and its association with oxidative stress, we assessed mitochondrial membrane potential in both control and PFOS-microinjected mouse oocytes using JC-1 staining because elevated membrane potential is often associated with increased ROS production. JC-1 is a lipophilic dye that naturally emits green fluorescence. Upon its accumulation within mitochondria at high concentrations, JC-1 forms reversible aggregates (J-aggregates), which emit red fluorescence. The red-to-green fluorescence ratio of JC-1 serves as a measure of mitochondrial polarization, with an increased red signal reflecting higher mitochondrial membrane potential.

Our results showed that the red fluorescence intensity of JC-1 staining in PFOS-microinjected oocytes was notably brighter than that in the control oocytes (Fig. 3A). Quantitative analysis showed no significant difference in the red-to-green fluorescence ratio between 2.4 nM PFOS-injected and control groups (Fig. 3B). However, the red-to-green ratio was significantly (*P* < 0.01) higher in the 5.6 nM PFOS-injected group compared to the control group (Fig. 3B), indicating that PFOS microinjection at this concentration elevated mitochondrial potential in oocytes.

**Figure 3.**
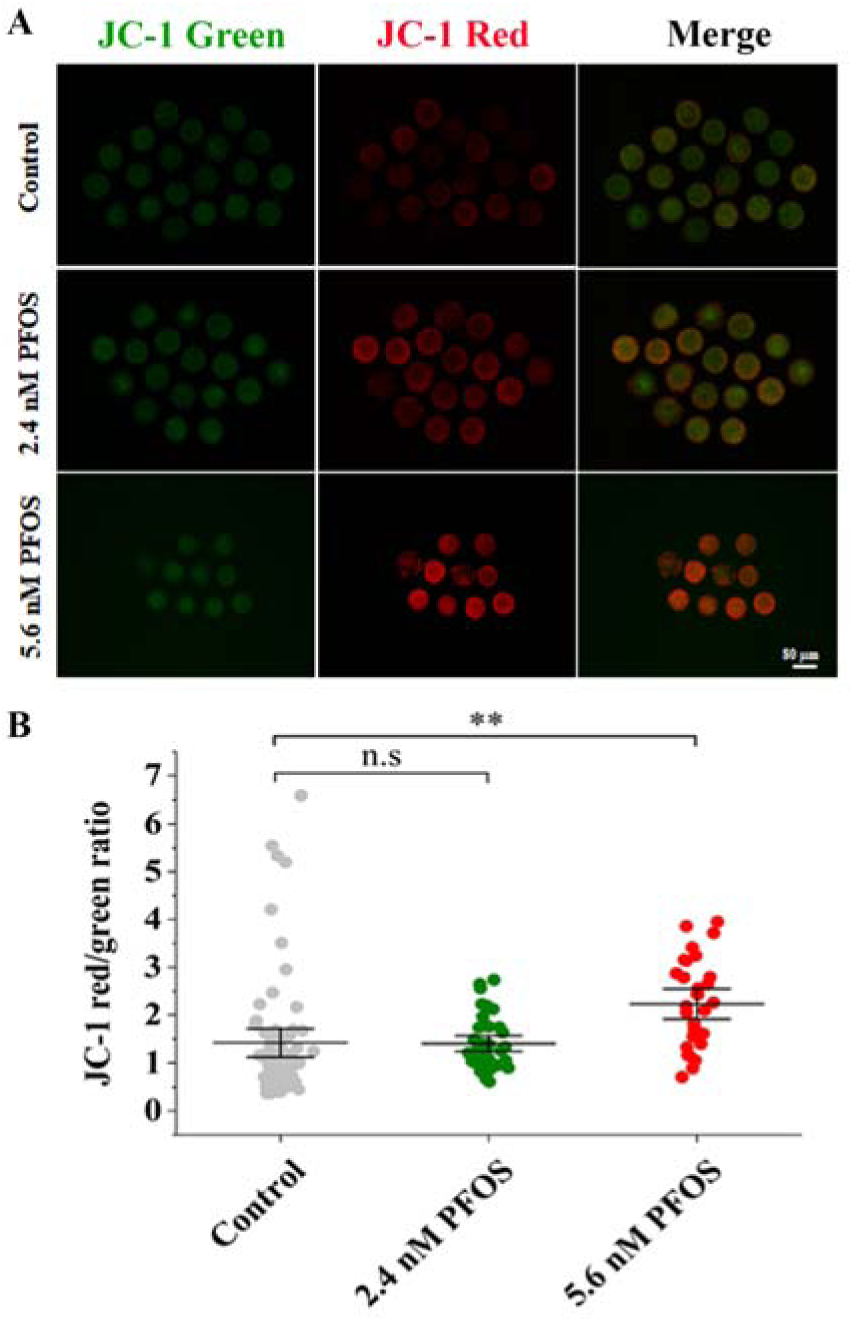
PFOS exposure increased mitochondrial membrane potential of 5.6 nM PFOS-injected oocytes. (A) Representative images of JC-1 staining, an indicator of mitochondrial membrane potential, in water-injected control, 2.4 nM, and 5.6 nM PFOS-injected oocytes. Scale bar: 80 μm. (B) Quantitative analysis of the JC1-red/green fluorescence intensity ratio in control, 2.4 nM, and 5.6 nM PFOS-injected oocytes. A total of 40 oocytes in the control, 40 in the 2.4 nM, and 31 in the 5.6 nM PFOS-injected groups were analyzed. Data are presented as mean ± SEM from at least three independent experiments. t-test, \*\**P* < 0.01 compared to the control.

### 3.4 PFOS microinjection induces chromosome misalignment and spindle abnormality in metaphase I (MI) oocytes

Since high ROS levels can disrupt oocyte spindle assembly and function, we hypothesized that PFOS exposure may result in abnormal spindle formation and misaligned chromosomes at MI, ultimately contributing lower PBE rates. To investigate this, we examined MI oocytes with or without PFOS microinjection (2.4 nM and 5.6 nM) using anti-α-tubulin immunostaining and DAPI counterstaining (Fig. 4A). Remarkably, the proportion of chromosome misalignment in MI oocytes was significantly higher in the 5.6 nM PFOS-injected group than the control group (*P* < 0.01, Figure S2). However, no significant difference was found between the 2.4 nM PFOS-injected and control groups. Next, we measured the spindle length and width, and MI-plate width of oocytes in control, 2.4 nM and 5.6 nM PFOS-injected groups (Fig. 4B). Spindle length was not significantly affected by PFOS microinjection (Figure S3). However, the spindle width was significantly reduced in the 5.6 nM PFOS-injected group compared to the control (*P* < 0.05, Fig. 4C); whereas no significant difference was detected in the 2.4 nM PFOS-injected group. The misalignment of meiotic chromosomes was observed after we measured the MI-plate width. We found that both 2.4 nM and 5.6 nM PFOS microinjections induced a significant increase in MI-plate width compared to the control (*P* < 0.05 and *P* < 0.001, respectively; Fig. 4D), indicating PFOS exposure disrupts chromosome alignment and segregation.

**Figure 4.**
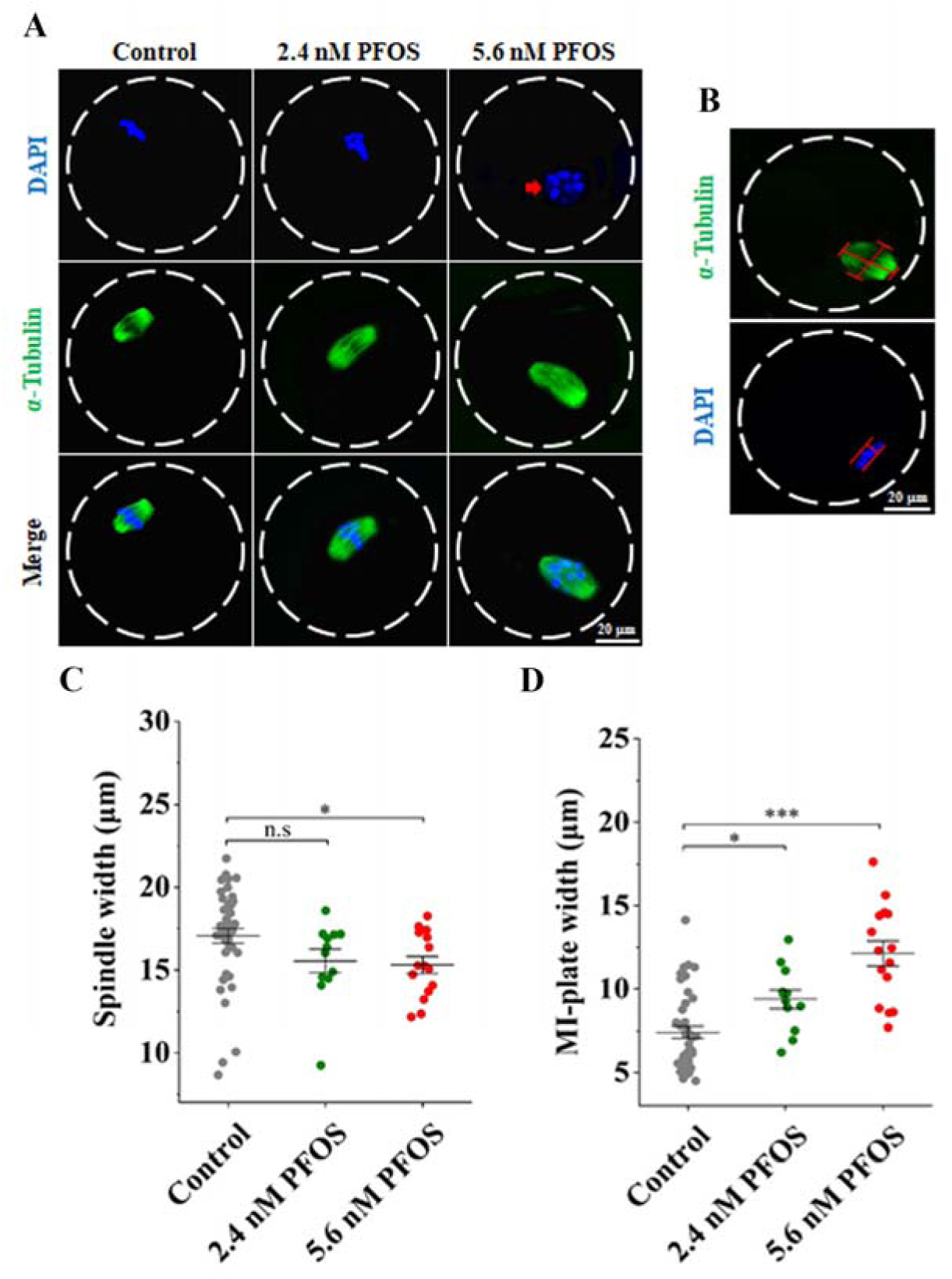
Effects of PFOS microinjection on metaphase I chromosome alignment in mouse oocytes. (A) Representative images of chromosome morphology after 6 hours of *in vitro* culture in the water-injected control, 2.4 nM, and 5.6 nM PFOS-injected oocytes. Red arrows highlight misaligned chromosomes. Scale bar: 20 μm. (B) Representative image of metaphase I (MI) spindle and chromosome morphologies in a control oocyte. Spindle length was determined by measuring the distance between the two spindle poles, while spindle width was assessed based on the microtubule width at the MI plate. The MI plate width was defined as the distance between the two outer edges of the DNA. α-tubulin is shown in green, and chromosomes/DNA are depicted in blue. Scale bar: 20 μm. (C) Quantitative analysis (mean ± SEM) of meiotic spindle width among the control, 2.4 nM, and 5.6 nM PFOS-injected groups. (D) Quantitative analysis (mean ± SEM) of MI plate width in the control, 2.4 nM, and 5.6 nM PFOS-injected groups. A total of 50 oocytes in the control, 22 in the 2.4 nM, and 16 in the 5.6 nM PFOS-injected groups were analyzed for chromosome misalignment and spindle morphology. Data are presented as mean ± SEM of at least three independent experiments. t-test, \*\*\**P* < 0.001, and \**P* < 0.05, compared with the control.

### 3.5 PFOS microinjection disrupts first polar body formation and delays meiosis II progression

Since PFOS microinjection affected MI progression, we next examined whether it also caused abnormalities at metaphase II (MII). After 24 hours of *in vitro* culture, most control oocytes reached the MII, whereas the proportion of oocytes arrested at prometaphase II was significantly higher in the 5.6 nM PFOS-injected group (*P* < 0.05, Fig. 5A), indicating that PFOS microinjection delays oogenesis. No significant difference was observed between the 2.4 nM PFOS-injected and control groups.

**Figure 5.**
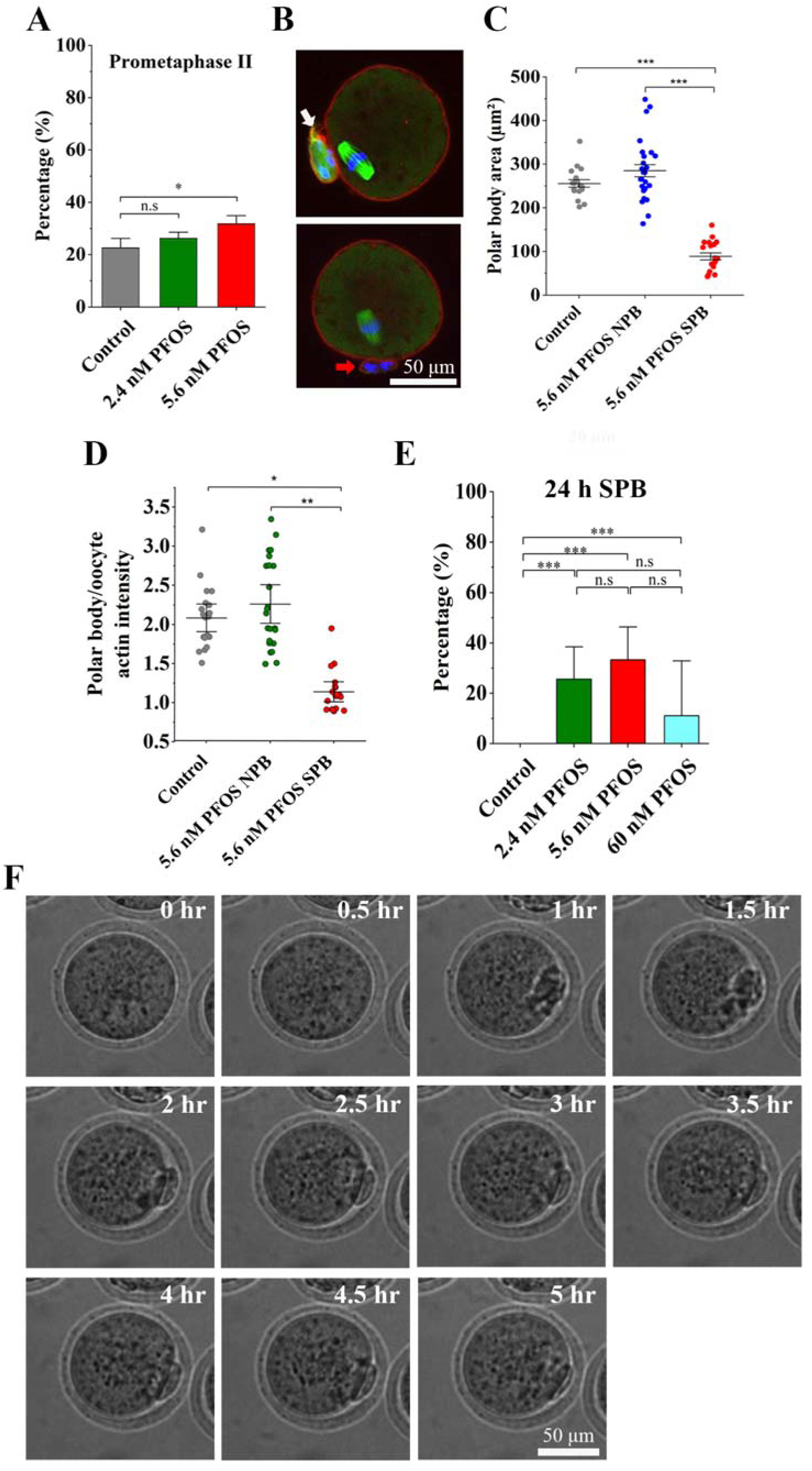
Effects of PFOS treatment on first polar body formation and meiosis II progression. (A) Percentage of oocytes arrested at prometaphase II in control, 2.4 nM, and 5.6 nM PFOS-injected oocytes after 24 hours of *in vitro* culture. (B) Representative images of polar body morphology in control (top image) and PFOS-injected (bottom image) oocytes after 24 hours of *in vitro* culture. Spindle in green, Actin in red, and DAPI in blue. The white arrow indicates a normal polar body (NPB) with typical morphology and visible spindles. The red arrow highlights a small polar body (SPB) lacking visible spindle structures. Scale bar: 50 μm. (C) Quantitative analysis (mean ± SEM) of polar body area among control, 5.6 nM PFOS-NPB, and 5.6 nM PFOS-SPB groups. (D) Actin intensity ratio (polar body/oocyte) in control, 5.6 nM PFOS-NPB, and 5.6 nM PFOS-SPB groups. (E) Percentage of oocytes extruding SPB in control and PFOS-injected groups (2.4 nM, 5.6 nM, and 60 nM) after 24-hour culture. (F) Time-lapse imaging showing the dynamics of small polar body formation following PFOS exposure. Data were presented as mean ± SEM from at least three independent experiments. t-test, \**P* < 0.05, \*\**P* < 0.01, \*\*\**P* < 0.001 vs. control. Scale bar: 50 μm.

Interestingly, PFOS-injected MII oocytes frequently exhibited unusually small polar bodies. To evaluate of PFOS-induced defects in polar-body morphology and size, we performed immunofluorescence staining of spindle microtubules (green), chromosomes/DNA (blue), and actin (red). In both the control and 5.6 nM PFOS-treated oocytes, normal polar bodies contained visible spindles (Fig. 5B, white arrow). By contrast, a subset of PFOS-treated oocytes displayed small polar bodies lacking spindle structures and appeared morphologically abnormal (Fig. 5B, red arrow). Quantification revealed that polar bodies from the PFOS-treated oocytes were significantly smaller than those in the control group and smaller than PFOS-treated oocytes that formed normal polar bodies (\*\**P* <0.001, Fig. 5C), further confirming that PFOS induces morphological abnormalities of first polar bodies.

To determine whether PFOS alters cytoskeletal organization, we quantified the relative actin intensity between the polar body and the oocyte. Oocytes with normal polar bodies in both the control and 5.6 nM PFOS-treated groups exhibited significantly higher polar body-to-oocyte actin intensity ratios compared to the 5.6 nM PFOS small polar-body group (*P* < 0.05 and *P* <0.01, respectively, Fig. 5D). This finding suggests that PFOS perturbs actin dynamics in a subpopulation-specific manner: normal polar-body extrusion is associated with robust actin accumulation, while small polar-body formation reflects impaired actin distribution. Consistent with this finding, microinjection of PFOS at all three concentrations tested (2.4 nM, 5.6 nM, and 60 nM) significantly (*P* < 0.001) impaired polar body extrusion, leading to increased small polar-body formation after 24 hours of culture (Fig. 5E). In contrast, injection of a high concentration of PFOS (60 µM) did not produce small polar bodies (n = 30), indicating that small polar-body formation is a low-dose-dependent phenomenon.

To further investigate the mechanism of small polar body formation, we performed live-cell imaging of PFOS-exposed oocytes. In oocytes with a small polar body, extrusion began approximately 1 h after culture initiation and the polar body enlarged until ∼3.5 h. Thereafter, the polar body progressive shrank, with pronounced size reduction observed by 5 hours (Fig. 5F). These dynamic observations indicate that PFOS compromises the stability of the extruded polar body, leading to abnormal shrinkage.

### 3.6 Embryonic development fail in oocytes with PFOS-induced small polar bodies

To evaluate the developmental competence of oocytes with PFOS-induced small polar bodies, we performed IVF on three groups: control, 5.6 nM PFOS with normal polar body extrusion, and 5.6 nM PFOS with small polar body extrusion. Many oocytes from both the control and the 5.6 nM PFOS-normal polar body group progressed beyond the 2-cell stage, whereas all oocytes from the 5.6 nM PFOS-small polar body group arrested before the 2-cell stage (Figure 6A). The proportion of oocytes arrested before the 2-cell stage was significantly higher in both 5.6 nM PFOS-treated groups compared with the control group (*P* < 0.01) (Figure 6B), indicating that PFOS microinjection impairs early embryo development. Notably, oocytes with PFOS-induced small polar bodies never reach the 2-cell stage, suggesting that defects in polar body extrusion directly compromise subsequent embryo development.

**Figure 6.**
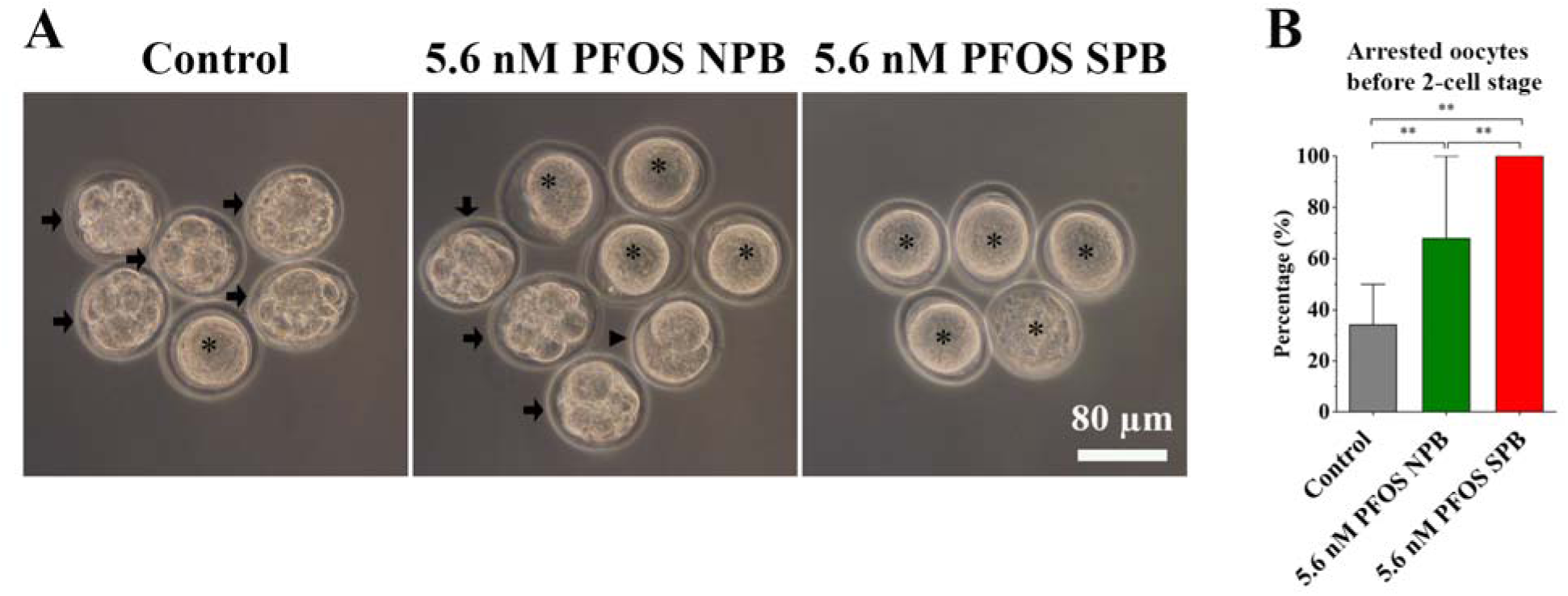
Effects of PFOS microinjection on mouse early embryo development. (A) Representative images showing embryonic morphology 48 h after *in vitro* fertilization (IVF) in the control group, the 5.6 nM PFOS-normal polar body group, and the 5.6 nM PFOS-small polar body group. Black arrows highlight embryos that developed beyond the 2-cell stage. The arrowheads point out the 2-cell embryos. The asterisks mark oocytes arrested before the 2-cell stage. Scale bar: 80 μm. (B) Proportion of oocytes arrested before the 2-cell stage in the control, 5.6 nM PFOS-normal polar body, and 5.6 nM PFOS-small polar body groups after 48 hours of IVF. A total of 38 oocytes (control), 36 oocytes (5.6 nM PFOS-normal polar body), and 28 oocytes (5.6 nM PFOS-small polar body) were analyzed. Data are presented as mean ± SEM from at least three independent experiments. t-test, **P < 0.01 compared with control.

### 3.7 Transcriptional differences between PFOS-injected oocytes with and without polar-body extrusion defects revealed by single-cell RNA sequencing

To investigate the transcriptional changes associated with small polar body (SPB) extrusion, we performed single-cell RNA sequencing on the PFOS-injected oocytes exhibiting either normal or small polar body phenotypes. Differential gene expression analysis identified 31 genes significantly upregulated and 50 genes downregulated in SPB oocytes compared to those with normal polar bodies (NPBs) (Fig. 7A). A hierarchical clustering heatmap demonstrated distinct expression profiles that clearly separated the two groups, with samples clustering according to polar body morphology (Fig. 7B). Genes in the upper and lower clusters displayed reciprocal expression patterns—upregulated in SPB oocytes and downregulated in NPB control oocytes, and vice versa—indicating that PFOS exposure alters gene expression in a manner dependent on the polar body morphology.

**Figure 7.**
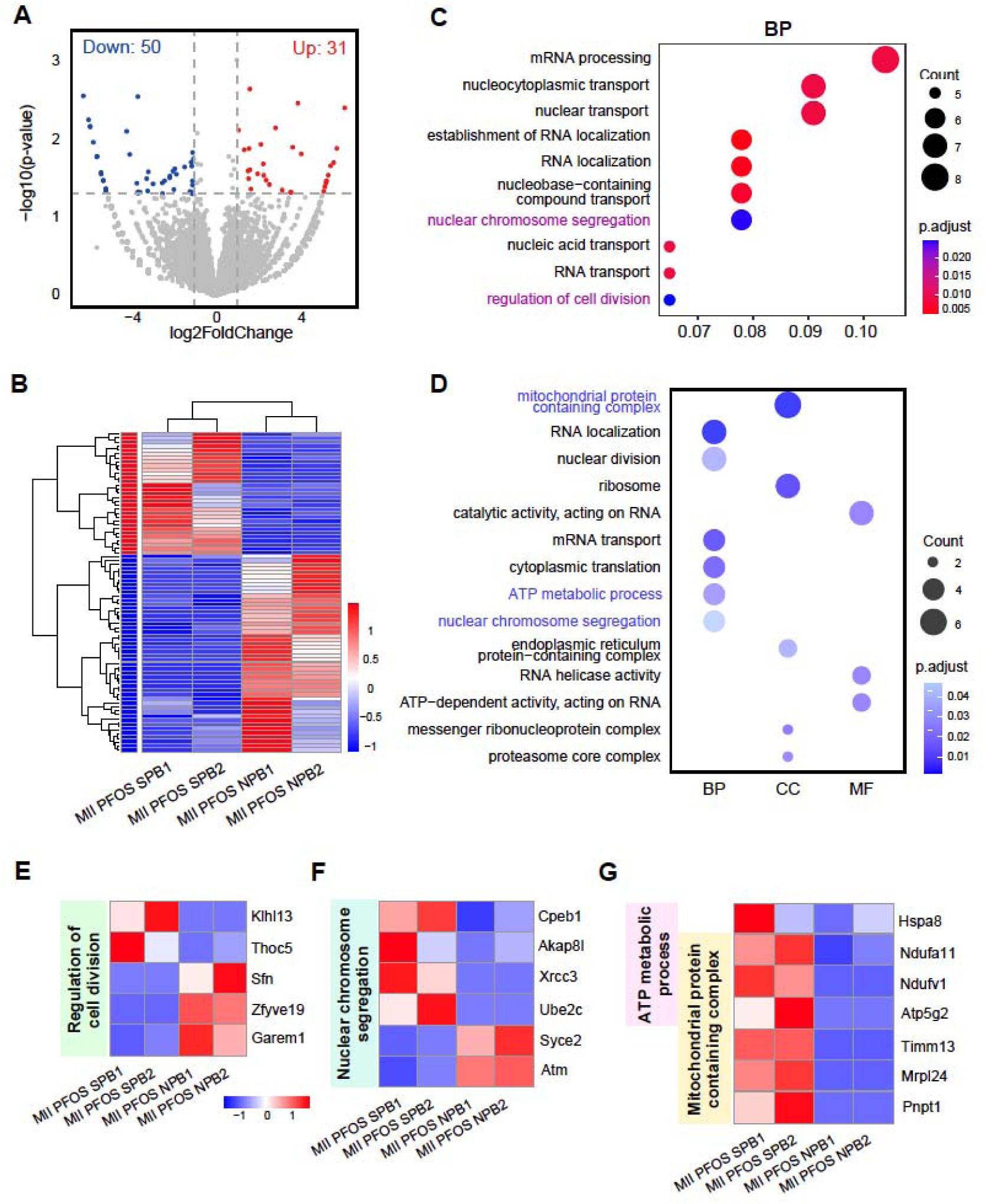
Transcriptomic comparison between PFOS-injected oocytes with normal and defective polar-body extrusion revealed by single-cell RNA sequencing analysis. (A) Volcano Plot showing differentially expressed genes (DEGs) between PFOS-injected MII oocytes with normal polar bodies (NPB, control) and those with small polar-bodies (SPB). Significantly upregulated genes (logLFoldChange > 1 and p-value < 0.05) are shown in red (Up: 31), and significantly downregulated genes are shown in blue (Down: 50). (B) Heatmap of DEGs across biological replicates (n = 2 per condition). Hierarchical clustering based on expression profiles shows distinct separation between MII oocytes with normal and defective polar-body extrusion. (C) Gene Ontology (GO) enrichment analysis – Biological Processes (BP) terms – revealing significant enrichment in processes such as regulation of cell division and nuclear chromosome segregation. Dot color indicates adjusted p-value; dot size reflects the number of genes enriched in each category. (D) GO enrichment analysis of 50 down-regulated genes (Fig 7A), showing significant enrichment in mitochondrial protein containing complex, ATP metabolic processes, and nuclear chromosome segregation across biological process (BP), cellular component (CC), and molecular function (MF) categories. Dot size represents gene count, and dot color indicates adjusted p-value. (E) DEGs involved in the regulation of cell division from panel C. (F) DEGs associated with nuclear chromosome segregation from panel C. (G) DEGs associated with ATP metabolic processes and mitochondrial protein containing complexes.

Gene Ontology (GO) enrichment analysis of the differentially expressed genes in the SPB group highlighted significant enrichment in biological processes such as mRNA processing, nucleocytoplasmic transport, and RNA localization (Fig. 7C). Importantly, terms such as “regulation of cell division” (Fig. 7E) and “nuclear chromosome segregation” (Fig. 7F) were also enriched, suggesting polar body extrusion is disrupted. Cellular component (CC) analysis revealed upregulation of genes associated with mitochondrial and ribosomal complexes, proteasomes, and cytoplasmic stress granules (Fig. 7D), pointing to mitochondrial dysfunction (Fig. 7G) and impaired protein homeostasis.

Collectively, these findings indicate that PFOS exposure triggers widespread transcriptional reprogramming in oocytes with small polar bodies, affecting pathways related to chromosome segregation and protein homeostasis. Such alterations may contribute to their abnormal morphology and impaired developmental competence.

## Discussion

This study provides compelling evidence that PFOS microinjection–even at physiologically relevant concentrations–impairs mouse oocyte maturation by disrupting meiotic progression, altering spindle architecture, increasing oxidative stress, and compromising mitochondrial function. These findings reveal novel mechanistic insights into PFOS-induced reproductive toxicity and extend observations from previous PFOS studies using high-dose culture-media supplementation (*17*, *27*, *28*).

Mammalian oocyte maturation is a highly intricate process that encompasses both tightly coordinated nuclear and cytoplasmic events that are essential for fertilization and embryogenesis (*29*). During the first meiotic division of oocytes, germinal vesicle breakdown exposes chromosomes to the cytoplasm, while spindle forms around them–rendering both structures vulnerable to chemical disruption (*30*). We observed that PFOS microinjection at follicular fluid (5.6 nM), and occupational (60nM) relevant levels significantly reduced GVBD and PBE rates, indicating a disruption in meiotic progression. Although several studies using culture medium supplementation have investigated the detrimental effects of PFOS on oocytes, the concentrations applied were relatively high, including 600 μM in mouse oocytes (*31*), 106 μM in bovine oocytes (*32*), and 800 μM in porcine oocytes (*33*). It has been found that PFOS exposure impedes meiosis, as evidenced by delayed spindle migration and sustained activation of the spindle assembly checkpoint (SAC) (*31*). Persistent activation of the SAC during metaphase I can lead to kinetochore-microtubule (K-M) attachment defects, chromosome segregation errors, and aneuploidy (*34*). PFOS interfered with the stability of K-M attachment and increased aneuploid rates (*31*). Thus, we speculated that PFOS microinjection, even at physiologically relevant levels, might disrupt kinetochore-microtubule (K-M) attachment, thereby compromise meiotic fidelity.

Accurate chromosome segregation in oocytes requires proper chromosome alignment and spindle assembly. Disruptions in spindle morphology or increases in metaphase I (MI) plate width can lead to chromosome misalignment, spindle instability, and increased aneuploidy risk (*22*, *35*). Our results showed that oocytes exposed to 5.6 nM PFOS exhibited prominent MI defects, including chromosome misalignment and increased MI-plate width, despite unchanged spindle length. These abnormalities, along with delayed progression to metaphase II (MII) and the absence of well-defined spindles in defective small polar bodies, indicate substantial disruption of spindle formation and asymmetric division.

Previous studies have reported that PFOS exposure disrupt actin dynamics by altering the localization of actin-related protein 3 (Arp3) and Palladin, two critical regulators of branched and bundled actin networks (*36*). Disruption of Arp2/3 complex or F-actin stability impedes spindle migration, asymmetric division, and successful oocyte cytokinesis (*37*). TimeLlapse confocal microscopy has confirmed PFOS-induced spindle migration failure in mouse oocytes (*17*). Taken together, the association of PFOS with actin disorganization and spindle migration failure strongly supports the conclusion that these mechanisms underlie the observed chromosome misalignment and defective small polar body formation in our experimental model.

Oxidative stress, characterized by an imbalance between reactive oxygen species (ROS) production and antioxidant defenses, is another established contributor to impaired oocyte maturation (*38*). Elevated ROS levels can lead to mitochondrial dysfunction, DNA damage, and meiotic spindle abnormalities, ultimately affecting female fertility (*39*, *40*). Mechanistically, PFOS-induced oxidative stress appears to be a central contributor to oocyte dysfunction in our study. PFOS exposure at 2.4 nM and 5.6 nM significantly increased ROS levels, as shown by elevated DCF fluorescence. Mitochondrial membrane potential was significantly increased in the 5.6 nM PFOS group, suggesting hyperpolarization. While this might initially reflect heightened mitochondrial activity, excessive mitochondrial potential has been linked to ROS overproduction and subsequent mitochondrial exhaustion or apoptosis (*41*). Consistent with previous reports (*17*, *28*, *31*, *42*, *43*), our results support that PFOS disrupts mitochondrial function and induces apoptosis, contributing to the high rate of degenerated oocytes observed in the 60 nM PFOS group.

Functionally, we found that 5.6 nM PFOS markedly impaired polar-body extrusion, and those oocytes exhibiting defective small polar bodies arrested before 2-cell stage following IVF. In contrast, No-PFOS control oocytes remained competent, PFOS-exposed oocytes with normal polar bodies also showed reduced developmental potential, highlighting the importance of polar body integrity. Previous studies have shown that high-dose PFOS significantly reduce polar body extrusion, induce aberrant symmetric cell division, increase aneuploidy risk, and impair early embryo development (*32*, *44*, *45*). Clinically, fragmented and morphological abnormalities in polar bodies have been correlated with diminished fertilization and impaired embryonic development (*27*, *46*, *47*). These studies suggest that PFOS impairs both the timing and quality of embryonic divisions. Our observation that oocytes with small polar bodies arrest before 2-cell stage are consistent with these studies, possibly due to PFOS-induced early cytoplasmic or structural defects during meiosis.

To further investigate the molecular basis of these defects, we performed single-cell RNA sequencing and identified robust transcriptomic differences between oocytes with normal and small polar bodies. We identified 31 upregulated and 50 downregulated genes in defective polar-body oocytes. Gene ontology (GO) analysis showed enrichment for terms related to “nuclear chromosome segregation” and “regulation of cell division,” indicating that PFOS exposure compromises the fidelity of meiotic division. These findings align with previous reports showing that PFAS can induce meiotic defects, including spindle abnormalities, chromosome misalignment, and oogenesis failure (*17*, *48*). In addition, small polar-body oocytes exhibited enrichment of transcripts associated with mitochondrial and ribosomal complexes, proteasome-related structures, and cytoplasmic stress granules (Fig. 7). Stress granules (SGs), which sequester mRNAs in response to cellular stress, play crucial roles in mitigating oxidative stress (*49*, 50). Their enrichment suggests that PFOS-exposed oocytes experience substantial cellular stress and activate compensatory response to PFOS-induced oxidative stress.

Together, these results strongly suggest that PFOS exerts its toxic effects through multifactorial mechanisms: impairing spindle assembly, inducing oxidative stress, disrupting chromosome alignment, and altering mitochondrial function. Notably, even low, environmentally relevant concentrations (2.4 nM) caused oxidative stress and polar body defects, raising serious concerns about low-dose PFOS exposure during oocyte maturation. Given the widespread presence of PFOS in biological and environmental compartments, our findings highlight the potential reproductive risk associated with chronic low-dose PFOS exposure. Future studies should explore downstream developmental consequences, including fertilization and embryonic competence, and investigate possible protective interventions–such as antioxidant supplementation–to mitigate PFOS-induced reproductive toxicology.

## Author contributions

Hasanur Alam and Huanyu Qiao conceived and designed the study. Hasanur Alam, Juan Dong, Vidhi Patel, and Wenjie Yang performed all bench experiments. Hasanur Alam, Shuangqi Wang, and Leyi Wang collected and analyzed the data. Hasanur Alam, Shuangqi Wang, and Huanyu Qiao wrote the paper. Hasanur Alam and Huanyu Qiao supervised the project. All authors read and approved the final manuscript.

## Acknowledgements

We sincerely thank all the members of Qiao Lab for their technical support and comments on this manuscript. This work was supported by NIH R00 HD082375, NIH R01 GM135549, and CCIL Seed Grant.

## Ethics

All animal experiments were approved by University of Illinois Urbana-Champaign (UIUC) Institutional Animal Care and Use Committee.

## Data availability

All data supporting this study are available within the article and its supplementary information. Single-cell RNA-seq data are deposited in the NCBI GEO (Gene Expression Omnibus) database with the accession number GSE315456. Additional datasets or related information are available from the corresponding authors upon reasonable request.

## Supporting Information

**Figure S1:** Schematic of the single-oocyte toxicological assessment workflow.

**Figure S2:** Additional analysis of chromosome alignment in MI oocytes. The percentage of chromosome misalignment was significantly higher in the 5.6 nM PFOS-injected group than the control group.

**Figure S3:** Additional analysis of MI spindle length in control and PFOS-injected oocytes.

