## Supplemental Figures for "Microinjection-based Single-Cell Toxicological Assessment Reveals How Physiological Levels of PFOS Impair Oocyte Maturation and Developmental Competence"

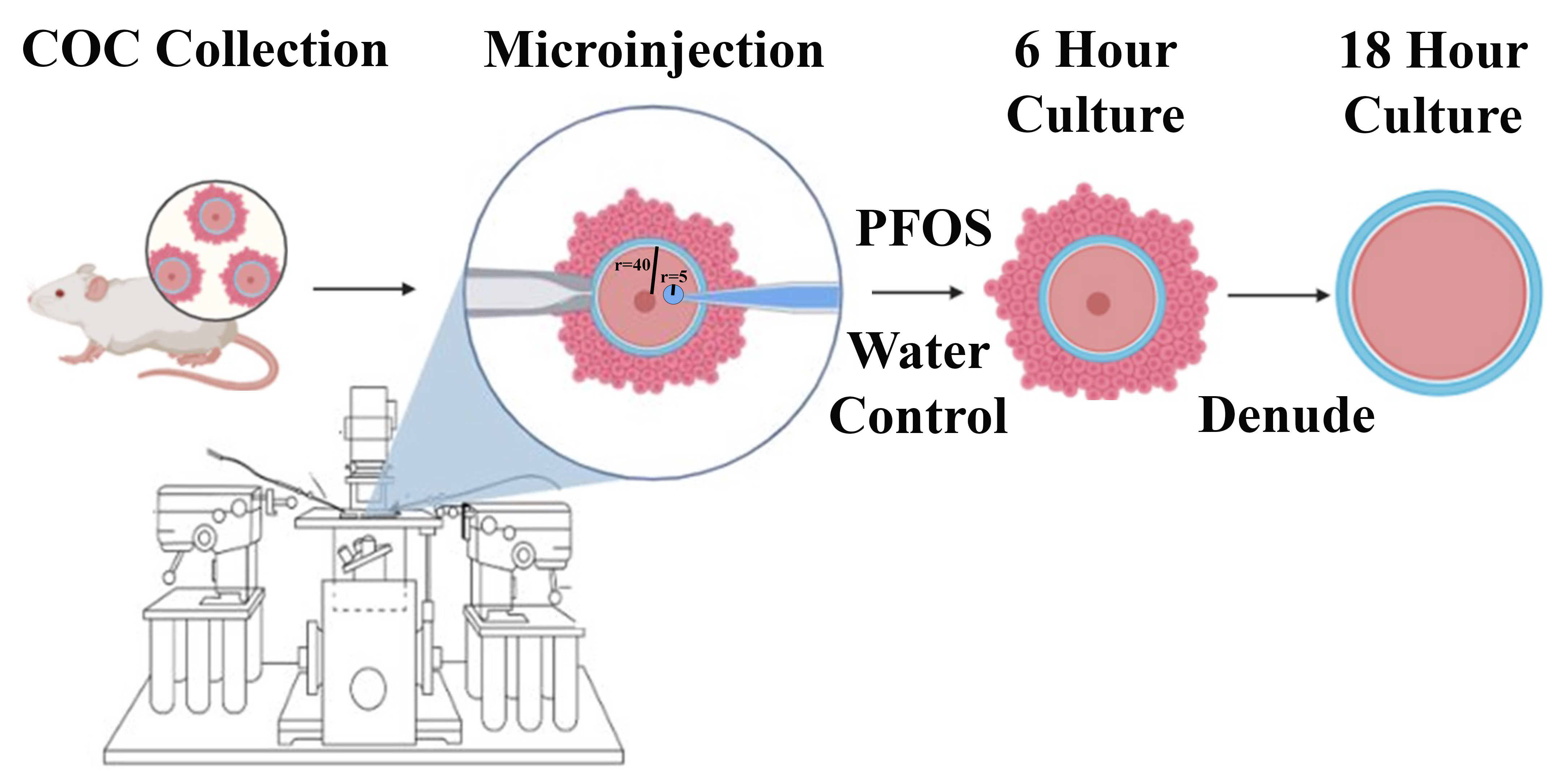

**Figure S1. Schematic of the single-oocyte toxicological assessment workflow.**

Cumulus oocyte complexes (COCs) were collected from mice for microinjection. The cumulus cells were removed from the oocytes after 6 hours of culture and then again cultured for 18 hrs. Microinjection of PFOS into oocytes enables precise control of the defined amount of PFOS delivered by bypassing the oocyte membrane. The volumes of both oocytes and PFOS droplets were calculated using the formula V = 4/3 πr³

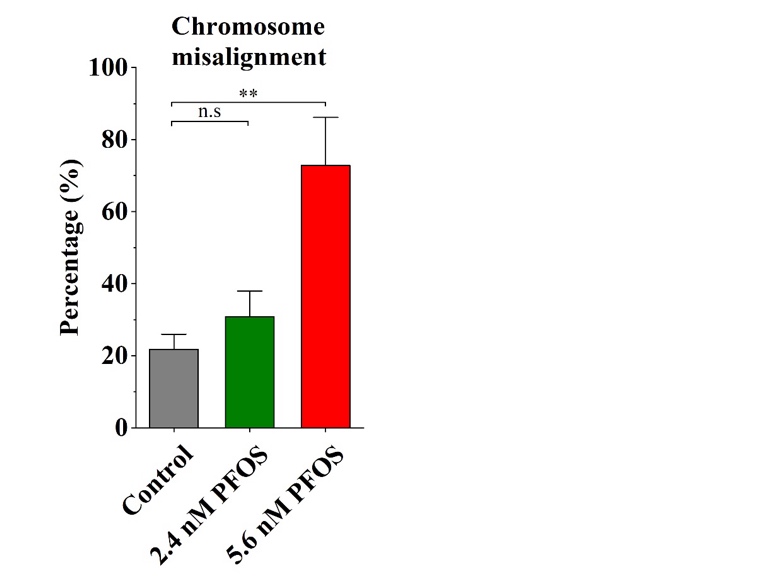

**Figure S2. Percentage of chromosome misalignment in control and PFOS-injected groups.**

Frequency of misaligned chromosomes in control, 2.4 nM, and 5.6 nM PFOS-injected groups. A total of 50 oocytes in the control group, 22 in the 2.4 nM PFOS-injected group, and 16 in the 5.6 nM PFOS-injected group were analyzed for chromosome misalignment. Data are presented as mean ± SEM of at least three independent experiments. Statistical significance was determined by t-test; **P < 0.01 compared with the control.

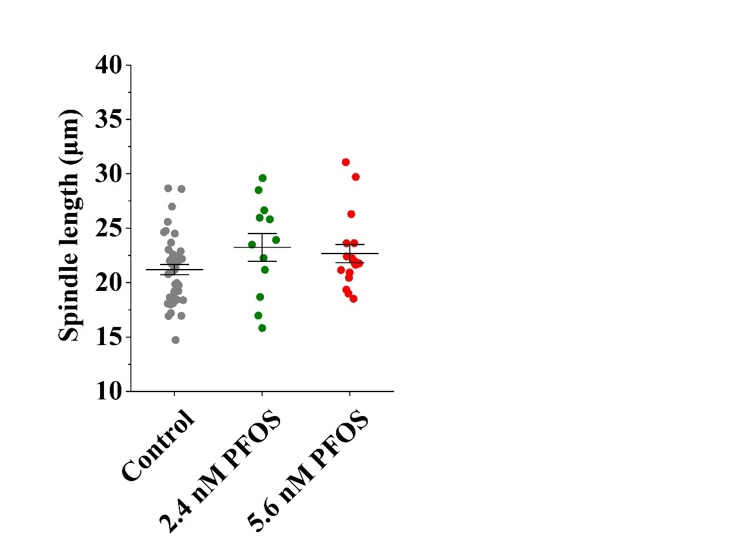

**Figure S3. Quantitative analysis of meiotic spindle length in control and PFOS-injected oocytes.**

Meiotic spindle length was measured in 50 oocytes from the control group, 22 oocytes from the 2.4 nM PFOS-injected group, and 16 oocytes from the 5.6 nM PFOS-injected group. Data are presented as mean ± SEM from at least three independent experiments. No statistically significant differences were detected among the groups by t-test.
